# OFC-SV: Interpreting structural variants in African and Asian cohorts with non-syndromic orofacial clefts

**DOI:** 10.64898/2026.10.13.757087

**Authors:** Kevin Chen, Yeonsoo Chung, Timothy Becker, Dashzeveg Bayarsaihan, Dong-Guk Shin

## Abstract

**Summary:** OFC-SV is a web-based data portal designed to help interpret structural variants in non-syndromic orofacial clefts genomic data published by The Gabriella Miller Kids First Pediatric Research Program. The portal offers a user-friendly way to explore 41 novel SV genes (of these 13 sharing regulatory perturbations already reported in the literature as SNV genes) potentially responsible for non-syndromic OFC from a cohort of children of African and Asian ancestry. A salient feature of the portal is its unique ability to capture epigenomic disruptions and complex oligogenic interactions in visualized regulatory pathways, thus advancing the frontier of disease etiology through convenient translation of computational signals into biological interpretations.

**Availability and Implementation:** OFC-SV is implemented in Python Dash as a web-based toolbox designed to help scientists review potential disease causal structural variants in non-syndromic orofacial clefts. This tool is freely accessible at: https://OFC-SV.cse.uconn.edu/ The source code and extra materials for this service can be downloaded from: https://github.com/KevinChen37/ofv-svexplorer

## Introduction

Orofacial clefts (OFCs) are one of the most common structural craniofacial birth defects globally affecting about 1.4 per 1000 births. The Gabriella Miller Kids First Pediatric Research Program has assembled a large-scale OFC related human genomics dataset, providing a unique opportunity to investigate how genomic variation contributes to OFC etiology. We carried out a pilot SV analysis focusing on non-syndromic orofacial clefts (nsOFCs)—cleft lip, cleft lip with or without cleft palate (CL/P), and cleft palate only—using a cohort of children of African and Asian ancestry made up of 241 parent–child trios Mukhopadhyay et al. (2020). To remove background SV signals, we used a large cohort of about 3,000 healthy genomes from the 1000 Genome Phase 4 (*G1KP4*) composed of 26 different ethnic populations Byrska-Bishop et al. (2022). OFC-SV is a data portal that enables OFC researchers to explore our findings from this SV focused analysis for nsOFC.

Motivation for developing OFC-SV is many fold. First, existing data analysis efforts on this resource focused mostly on SNV analysis, and little attempt has been made to perform Structural Variant (SV) analysis. Second, the datasets required to study nsOFC are all scattered around in various primary data sources (e.g., Kids First data portal, ENCODE, Gene Ontology, FaceBase and others), and thus eliciting plausible clues to explain the disease etiology though an integrated analysis is extremely cumbersome. Third, even if the appropriate collection of related datasets is assembled and the integrated data analysis is performed, presenting the analysis outcomes in a way that ordinary OFC scientists to explore the findings is challenging because of a lack of a user-friendly way of examining the analysis outcomes in a public web portal.

Another important consideration for building the OFC-SV data portal is to address the dbGap controlled-data access compliance issue due to the sensitive nature of the personal genomics data. OFC-SV is limited to presenting Population view for the gathered samples and yet enable scientists to explore how epigenomic features may influence the disease etiology through complex oligogenic interactions.

## System design concept

Our SV analysis consists of two steps: SV calling, which identifies structural variants (SVs) in each sample, and evidence tracking, which gathers evidence to help explain the potential causal relationships underlying each sample’s phenotype. Additional details on the system design are provided in Sections A, B, and C of the Materials and Methods section of the Supplementary Material.

### Primary data sources

We downloaded the whole genome sequence (WGS) data for African and Asian ancestry from the Kids First Portal Mukhopadhyay et al. (2020). From the original dataset we excluded a few due to potential annotation issues (e.g., single parent, code mismatching, etc.) and used 241 parent–child trios for our SV analysis.

### SV Calling

The first step is applying our existing ensemble method, called FusorSV Becker et al. (2018) which employs a comprehensive set of machine learning algorithms to combine data from eight specialized SV callers Lam et al. (2010); Fan et al. (2014); Abyzov et al. (2011); Rausch et al. (2012); Layer et al. (2014); Chen et al. (2016); Cameron et al. (2021); Chen et al. (2014). It detects a wide range of SVs from 50 bp up to and beyond 100 Mb, including full chromosomal aneuploidies. The SV subtypes detected include deletion (DEL), insertion (INS), duplication (DUP), and inversion (INV) in the order of highest to lowest occurrence in human populations. This ensemble method uses at least two callers for each subtype, with every call having a 0.75 reciprocal overlap threshold. The SV clusters are then used to calculate allele frequencies in the background dataset (1000 Genome Project Phase 4; *G1KP4*) Byrska-Bishop et al. (2022). Allele frequencies below 5% in the background population are defined as rare, whereas SVs absent from the background dataset are assigned a frequency of 0% and considered de novo. In addition, we counted the number of samples from the OFC cohort (parents and children) to determine allele frequencies separately within the parental and affected-child groups.

### Evidence Tracking

The second step is associating each identified SV with a set of related genomic features exerting regulatory functions such as exons, promoters, enhancers and insulators. We used histone marks from the H9 cell line study from ENCODE (https://www.encodeproject.org) to determine candidate regulator evidence for each cell type, with our area of interest being Mesenchymal Stem Cell (MSC) and Neural Crest (NCC). We also utilized *H3K27me3* ChIP-seq peaks and transcription start sites (TSSs) from the RefSeq known gene annotation table for hg38 (UCSC GoldenPath) (https://hgdownload.cse.ucsc.edu/goldenpath/hg38/bigZips/). Candidate promoter regions were defined by capturing *H3K27me3* peaks that fell within a 3-kb window upstream of each TSS. We then used the promoter candidate areas for each cell type to further annotate enhancer regions (*H3K27ac* and *H3K27me1* for active and poised enhancers respectively). Insulator candidates were drawn from the CTCF peaks for each cell. To further investigate the regulatory nature of the genes in each area, we also explored the downstream evidence by way of examining the normal palate tissue gene expression levels in single-cell mouse palate tissue across the developmental time points where variations would be most disruptive to development Yan et al. (2024). We also examined the variant-to-gene mapping information (V4-hg38.Gene-Links.3D-chromatin.txt) to check if each disrupted enhancer has been annotated to distant genes through a 3D loop physical contact Moore et al. (2026). Lastly, our SV genes of interest are compared with OFC genes annotated in literature to produce a total of 41 novel SV genes and of these 13 sharing regulatory perturbations already reported in the literature as SNV genes (Supplement Table 1).

### Pathway modeling

One salient feature of OFC-SV is the curated regulatory network that helps scientists explore significance of identified SV genes within a biological pathway map. This pathway map focuses on modeling relationships among genes and two standardized GO terms, Molecular Functions (MF) and Biological Processes (BP) which constitute nodes in the regulatory network. These nodes are then interconnected with labels that convey the relationships generally known between two participating nodes. The example in Fig 1 illustrates how we are extending the well-established ns OFC regulatory knowledge *TP63 → IRF6 → GRHL3 → CLDN3* promoting epithelial/periderm differentiation and cell-cell adhesion by adding additional nodes and relationships. Our way to render the regulatory network was motivated not only to make the map easily understandable by scientists, but also by Large Language Models (LLM) modules that eventually should be able to exploit this type of resource. Note that here we introduced an edge labeled with “putatively-regulate” because our curation of SV gene mapping should be considered hypothetical based on an array of evidence.

**Figure 1.**
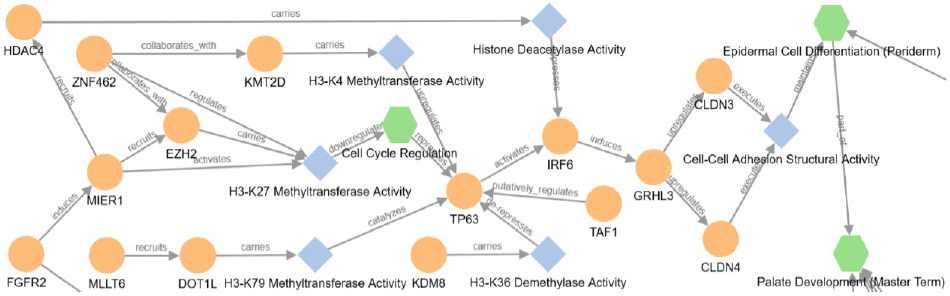
Extending the established *TP63-IRF6-GRHL3* axis

## Implementation

The developed OFC-SV is written in the Python programming language (version *≥* 3.6.0) and takes advantage of the Dash Dabbas (2021) for the pipelined and customizable visualization to produce publication-ready figures and tables. For sequence interval operations and pathway map creation, PyRanges and Cytoscape were used, respectively.

### Static and Interactive Data Visualization

The Data View tab of the portal provides various data visualization plots showing sample demographics and the distribution of discovered SVs. Some notable plots compare the distributions of the four major SV types across different cohorts. Fig 2a and 2b compare SV type distributions between African and Asian cohorts. The African cohort exhibits a slightly higher proportion of deletions (DELs; 75.5%) than the Asian cohort (73.7%). This trend is consistent with previous reports describing SV patterns across ancestral populations and phenotypic groups Ho et al. (2020).

**Figure 2.**
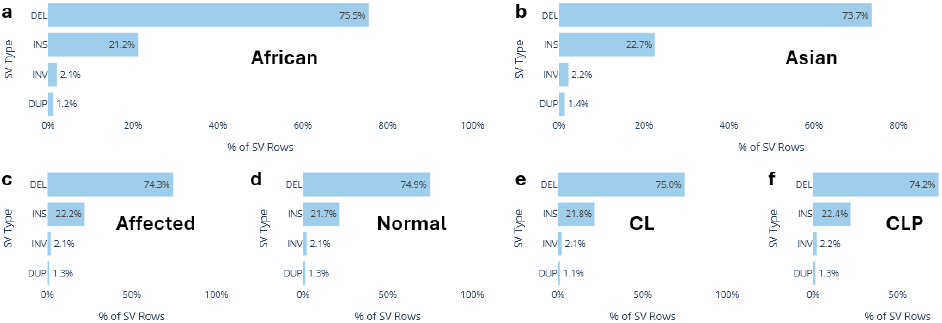
Distribution patterns of SV types across different sample groups

Fig 2c and 2d compare SV type distributions between Affected and Normal samples and reveal highly similar overall patterns. Likewise, Fig 2e and 2f demonstrate comparable distributions between the CL and CLP phenotype groups. The similarity observed across these comparisons suggests that disease relevance may depend less on the overall frequencies of SV classes and more on the specific combinations of SVs and the genes they affect. This interpretation is particularly supported by the near-identical distributions in Fig 2c and 2d despite the approximately 1:2 sample-size ratio between the Affected and Normal groups. Additional SV distributions are provided in Section D (Various Statistics) of the Supplementary Material.

The portal also provides an Interactive sub-tab that allows users to generate customized boxplots by selecting different X- and Y-axis variables. For example, by setting Chromosome and SV Type as X-axis variables and Unique SVs as Y-axis variable, users can identify unusual patterns such as chromosome 2 containing approximately three times more inversions (INVs) than duplications (DUPs) within a phenotype group.

### Case studies of various SV disruptions

To demonstrate how different types of SV disruption can be explored, we present six case study examples using the categorization scheme of “where the SV occurs” and “what regulatory element it disrupts”. Each of these cases can be explored in depth from the OFC-SV web portal. Additional case study details including extended pathway diagrams are provided in Section E (Pathway Modeling Case Studies) of the Supplementary Material.

#### Case Study 1. Coding exon disruption (TET3)

*TET3* encodes a DNA demethylation enzyme (Ten-Eleven-Translocation 3) and is one of the three mammalian TET proteins (*TET1/2/3*) that remove or modify DNA methylation marks. It functions as a master epigenetic regulator of developmental gene expression and acts directly on the epigenome to control activation of developmental genes. We found a deletion SV (DEL; C 16066, chr2: 74047046-74047999, size 954 bp) which lies entirely within Exon3 (chr2:74046278-74048411, size 2134 bp, freq:0.0042) of the gene. This SV is translated into an in-frame deletion of 318 amino acids, accounting deletion of about 45% of the exon, potentially resulting in removal of a functional domain or creating a hypomorphic protein. In addition, this deletion SV also overlaps over 95% with an active enhancer (chr2: 74047746-74048023, neural crest cells) which itself lies within the exon region. In the pathway viewer this gene is tied to DNA Demethylase Activity (GO:0035514) that activates TGFB3. As DNA Demethylase Activity is counter-balanced by DNA Methyltransferase Activity (GO:0003886), disruption in *TET3* can suggest an instability in the eventual Palate Development.

#### Case Study 2. Promoter/TSS disruption (KDM8)

*KDM8* is an epigenetic/chromatin regulator influencing chromatin state, cell proliferation, DNA damage responses, and microtubule/cytoskeletal organization. We found a deletion SV (DEL; C 148215, chr16: 27204873-27207194, size:2322 bp, freq:0.0042) that overlaps with *KDM8* ‘s promoter/TSS region identified for neural crest cells. Being an epigenetic regulator belonging to the JmjC-domain protein family, it regulates *alpha*-tubulin acetylation and microtubule stability and facilitates cytoskeletal remodeling associated with epithelial-to-mesenchymal transition. Noticeable from the OFC-SV’s pathway viewer is that *KDM8* and *DOT1L* function as opposing biochemical forces on the epigenome, one serving as an epigenetic eraser (demethylase) and the other as an epigenetic writer (methyltransferase), respectively. Both molecular functions drive the cellular plastic changes through the well-established *TP63-IRF6-GRJHL3* axis leading to Epidermal Cell Differentiation (Periderm) in palatogenesis, and the over-2-kb deletion near the *KDM8* promoter region can cause a significant disruption in this aspect of the balancing act.

#### Case Study 3. Intergenic distal enhancer disruption (MLLT6)

*MLLT6* (also called *AF17*) is a component of the *DOT1L* histone H3K79 methyltransferase complex. It binds/associates with *DOT1L* and helps enhance its methyltransferase activity Okuda et al. (2017). We identified two SVs that overlap distal enhancers located upstream and downstream of MLLT6, respectively: an insertion (INS; C 226473; chr17:38,751,016–38,751,335; 320 bp; freq: 0.0042) overlapping a distal enhancer 45.7 kb downstream of the MLLT6 transcription start site (TSS), and a deletion (DEL; C 154706; chr17:38,663,437–38,664,493; 1,057 bp; freq: 0.0042) overlapping a distal enhancer 40.8 kb upstream of the MLLT6 TSS. The INS SV overlaps with two No-cleft Embryo cCREs (EH38E4161022 and EH38E3221048) both of which are classified as pELS. ChIA-PET interactions from SCREEN for both cCREs include the identical set of targets including *MLLT6, EPOP, MED1* and several others. The pathway view illustrates how *MLLT6* is linked to H3-K79 Methyltransferase Activity through *DOT1L* and then to the well-established *TP63-IRF6-GRJHL3* axis leading to Epidermal Cell Differentiation (Periderm) in palatogenesis and eventually to EMT.

#### Case Study 4. Intron enhancer disruption (LMO7)

*LMO7* is a multi-domain scaffold protein playing key roles in cell adhesion, nuclear structure, cytoskeletal organization, and transcriptional regulation. It acts like a scaffold/linker that connects adhesion proteins to the actin cytoskeleton and has a reported role in organizing epithelial structures of embryonic ectoderm. We found a large deletion SV (DEL; C 130729, chr13: 75788946-75793975, size: 5030 bp, freq: 0.0042) in *LMO7* that entirely overlaps with multiple active enhancers in mesenchymal and a cluster of dELS type No-cleft Embryo cCREs (EH38E307178, EH38E307179 and EH38E3071711). The ChIA-PET link of all three enhancers in SCREEN is *LMO7*, suggesting a self-promoting enhancer. In the pathway view considering the biological functions (adhesion, cytoskeletal organization, epithelial integrity, EMT-related processes), *LMO7* is tied to Epithelial Cell-Cell Adhesion (GO:0090136) which is linked to Epithelial-Mesenchymal Transition (EMT) (GO:0001837). Considering the 5 kb deletion and its self-enhancer deletion suggested by multiple ChIA-PET links, this deletion suggests a strong biologically plausible regulatory effect.

#### Case Study 5. Intergenic insulator/CTCF disruption (TAF1)

*TAF1* is a master transcription-initiation regulator and is one of the core components of the *TFIID* complex that recruits RNA polymerase II to promoters and initiates. It acts as a global transcription machinery component of embryogenesis such as neurodevelopmental disorders, intellectual disability syndromes, and developmental abnormalities Crombie et al. (2024). We found a very large deletion (DEL; C 90036 ; chrx: 71314241–71352748, 38508 bp; freq: 0.0042) located in about 13.4 kb intergenic upstream of *TAF1*, that overlaps with CTCF/insulator cluster known in mesenchymal, H9 and other tissues. This DEL SV demonstrates an example of a paternal X inheritance affecting daughter. Also noticeable is the cluster of PLS type enhancers that are located near *TAF1* TSS and their targets are *TAF1* upstream genes such as *ZMYM3, NONO, GJB1, ITGB1BP2* and others. The SV overlapping insulator clusters can be the classical example of barrier designed to block expression of *TAF1* in the sense of Hi-C 3D TAD domain, and its disruption can be functional. *TAF1* being a core component of the *TFIID* complex, our pathway extension places *TAF1* to “putatively regulate” *TP63* and its subsequent *IRF6-GRJHL3* axis leading to EMT.

#### Case Study 6. Intronic insulator/CTCF disruption (PRKCA)

*PRKCA* encodes protein kinase C alpha, a signaling kinase involved in cellular proliferation, differentiation, adhesion, and cytoskeletal processes—all involved in craniofacial morphogenesis including mesenchymal proliferation/differentiation, migration, epithelial–mesenchymal interactions Ji et al. (2020). We found a deletion SV (DEL; C 156268; chr17: 66562075–66562164, 90 bp; freq: 0.0042) located in intron No. 3 of the gene and it overlaps an insulator candidate (chr17:66561968-66562967; 1000 bp; Cell: Mesenchymal; Score:5.0). Noticeable in this insulator overlapping deletion SV is the cluster of total four deletion SVs of similar sizes (C 156266, 100 bp; C 156268, 90 bp; C 156269, 110bp) overlapping with each other and all deleting the leading portion of the intron No. 3 insulator. Similar deletion SVs are also found in background with the frequency of 11 samples. Nevertheless, the frequency comparison shows a significance enrichment in affected samples, that is, 0.0168 (=4/238) in children vs. 0.0036 (=11/2981) in background 1000G samples. Interestingly, the paternal inheritance of an SV disrupting *DNAJC5* (DEL) and *MKKS* (DUP) was observed in a family of both father and daughter, whereas only the daughter carrying an additional *PRKCA* insulator DEL exhibiting the nsOFC phenotype. *MKKS* is involved in ATP binding and chaperone-related functions, while *DNAJC5* participates in cellular protein homeostasis, vesicle trafficking, and neuronal/synaptic processes. Together, these findings support a potential oligogenic model in which the combined disruption of *DNAJC5, MKKS*, and *PRKCA* contributes to nsOFC susceptibility, where the *PRKCA* insulator deletion potentially serves as a threshold-crossing event that determines phenotypic manifestation. In the pathway we place this gene as a parallel upstream signaling node that feeds an inhibitory edge directly into Sonic Hedgehog (SHH) node, balancing out the positive flux driven by the ciliary localization machinery, whereas *MKKS* and *DNAJC5* are dedicated structural helpers that build the ciliary factory.

### Use by the OFC research community

The program is available to the public as a web-based tool at https://OFC-SV.cse.uconn.edu. The SV analysis outcomes are limited to the datasets from the cohort of families of African and Asian ancestry. Exploring the processed data can begin by reviewing the summary statistics of Data Overview tab. Table Inspection tab enables the user to peruse the entire 41 SV genes that have been curated with evidence and can be used as a gateway to explore details of evidence and reasoning behind choosing the genes. Specifically, Gene and FusorSV id(s) column values are interactive and by clicking the column values will present a dropdown menu enabling the choices. Population tab shows IGV view specific to the chosen gene or SV of interest. It provides multiple configurable genomic annotation tracks to review evidence. Pathway tab enables users to review how one or more genes of interests are mapped on regulatory mechanisms leading to EMT in the context of palatogenesis. Lastly, Case Study tab offers more detailed explanation of the six SV disruption cases discussed in the above and the description is interlinked to conveniently explore related topics in IGV and Pathway map.

## Conclusions

OFC-SV Explorer is a publicly accessible, interactive web tool that enables biomedical scientists to investigate the potential involvement of structural variants (SVs) in nonsyndromic orofacial clefting (nsOFC). In contrast to broad community resources such as FaceBase, OFC-SV Explorer is a highly specialized platform dedicated to a single congenital disorder, offering focused analyses tailored to nsOFC research. Currently, the portal reports foundational SV analyses, illustrating how different classes of structural variants are distributed across the genome. Although the current dataset includes 241 trios of African and Asian ancestry, it provides an initial view of SV landscape patterns in nsOFC. Beyond descriptive analyses, the pathway map component extends the portal’s utility by integrating existing knowledge of nsOFC gene regulatory networks and highlighting how structural variants may perturb these pathways. This framework enables researchers to evaluate SV findings within a broader biological context and generate hypotheses regarding disease mechanisms. More importantly, the portal provides a foundation for developing oligogenic models of nsOFC in which the combined effects of multiple disrupted genes contribute to disease risk. Such models may help elucidate how concerted genetic perturbations converge on developmental pathways to produce the cleft phenotype, like the one we reported elsewhere Becker et al. (2026).

## Contributor Information

KC and YC implemented the portal. TB generated data. DB reviewed and DS wrote the manuscript.

## Supplementary data

Supplementary data are available online.

## Acknowledgments

This work was supported in part by the National Institutes of Health [NIDCR-5R03DE033083].

## Data Availability

All the example data and analytic code for this web service are available at https://github.com/KevinChen37/ofv-svexplorer. FusorSV is available at: https://github.com/timothyjamesbecker/Fu

## Supplementary Material

### Data downloads and processing pipelines

1. Downloaded CRAM files from dbGAP (via CAVACTICA bulk download command line tool): https://kidsfirstdrc.org/
2. Downloaded multiple phenotype files and generation of a harmonized phenotype JSON file using kfd join tsv pheno.py script.
3. Downloaded SNV family VCF files from KFD portal for use with ERDS.
4. Downloaded 3000 CRAM files from G1KP4 data portal: https://www.internationalgenome.org/data-portal/sample
5. CRAM to BAM file conversion (for SV calling) using samtools cram to bam.py script
6. Validation and BAM check using samtools index stats.py script.
7. Run batches of BAM files (out of 742 KFD and 3000 G1KP4 samples) on server with 8 SV calling algorithms (GRIDSS, MELT, Manta, Delly2, CNVpytor, ERDS, BreakDancer, LUMPY) using the kfd batch.sh and g1kp4 batch.sh scripts and (individual runner scripts in supplements)
8. Built a custom FusorSV fusion model to G1KP3 gold-standard data (as in the paper) for the 8 SV calling methods.
9. Applied the custom fusion model to the joint batch of KFD and G1KP4 VCF data (4K samples x 8 VCF files each 32KVCF files) to merge and filter calls to produce individual VCF files for each sample.
10. Downloaded HM ChIP-seq (H3K4me1, H3K4me3, H3K27ac, CTCF) data from ENCODE human H9 stem cell portal: (https://www.encodeproject.org)
11. Downloaded HiC data from ENCODE human motor neuron, cerebellum astrocyte human data (https://www.encodeproject.org)
12. Installed and used JASPAR DB for Transcription Factor binding site API to determine TFBSS enrichment in SV regions of interest.

## Materials and Methods

### A. Whole-genome sequencing (WGS) data

The Gabriella Miller Kids First Pediatric Research Program aims to uncover the genetic etiologies of pediatric cancers and congenital anomalies. By generating whole-genome sequencing (WGS) data from affected families, the program provides an open-access resource for community-driven scientific analysis. The datasets generated from this program include short-read and long-read WGS data (alongside whole-exome sequencing, RNA-Seq, and phenotypic/clinical data) for thousands of affected children and their family members (often in trios or multi-generational families). Regarding non-syndromic OFC datasets, it includes 30× Illumina paired-end WGS data along with demographic information based on genotype and phenotype identifiers from Asian, African, Latin American, Filipino, and European cohorts comprising parent–child trios. Our study has been limited to processing WGS data limited to trio cohorts from African and Asian populations, specifically 241 trios from Kids First and 2981 samples from the 1000 Genomes Phase 4 project.

### B. Structural Variant Data Processing

Our SV analysis follows a comprehensive and robust protocol comprising eight specialized SV callers, namely, BreakDancer, CNVpytor, LUMPY, DELLY, Manta, ERDS, MELT, and GRIDSS2, that power an ensemble called FusorSV. Using this system, we can detect a wide range of SVs from 50 bp to more than 100 Mbp, including full chromosomal aneuploidies. These include deletions, insertions, duplications, and inversions, in the order of highest to lowest occurrence in human populations. In general, the ensemble method uses at least two callers for each subtype, where MELT is specialized for insertion detection and CNVnator excels in discovery of large deletions and duplications. FusorSV uses previously validated SVs as part of its supervised learning framework to determine the best tool combinations and cutoffs for removing overlapping or low-confidence calls. Its merged results achieved an *in vivo* validation accuracy exceeding 80% as reported by Becker et al. (2018). We define “rare” as occurring in 1% or less of the G1KP4 population. We define de novo as both rare and absent from unaffected parents. Population frequency for each SV cluster is calculated as

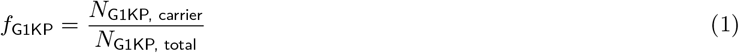

where *N*_G1KP,carrier_ is the number of population-background individuals carrying the corresponding SV and *N*_G1KP,total_ is the total number of background individuals analyzed. A frequency of zero therefore indicates that no corresponding SV satisfying the clustering criterion was observed in the population reference.

#### Eliminating background noise

We implemented a case-to-control filtering system that dynamically removes background noise using reference datasets. For our analysis, control data consist of Illumina paired-end sequencing from the 1000 Genomes Phase 4 dataset, comprising about 3,000 healthy samples. Once the filtering is completed, the noise eliminated data is grouped and merged into each trio using the VCF specification pedigree notation along with phenotype designations. The first phase of SV data processing is summarized in Figure S1.A, i.e., generating the initial group of novel SVs.

**Figure S1.**
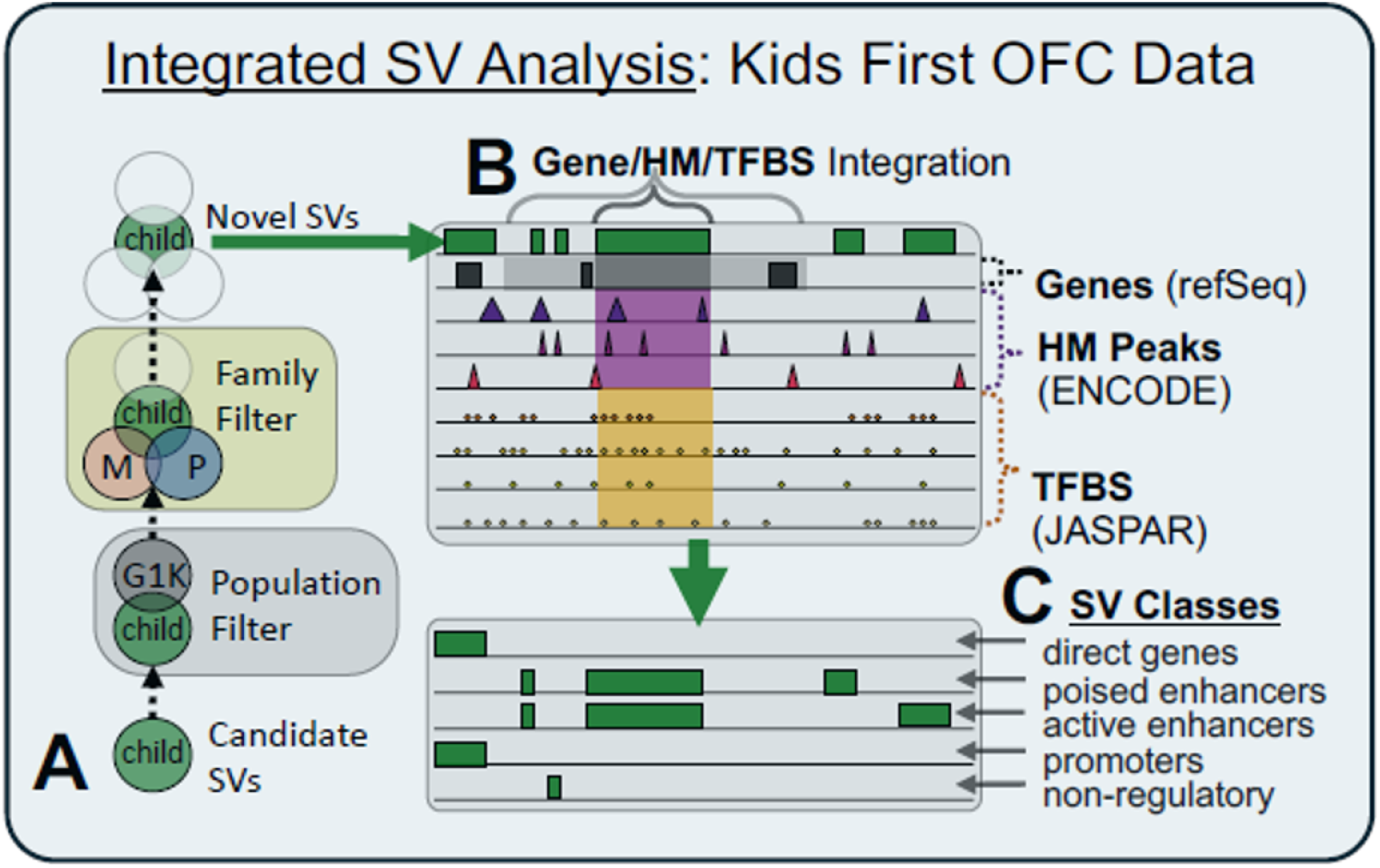
A. Generating novel SV candidates, B. Controlling false positives, C. Grouping and Classifying SVs.

#### Controlling false positives

Many SV callers produce numerous false positives. It is critical to employ a means to control false positives to generate high-confidence calls. We use a separate filtering method, called CellVar (Becker et al. (2022), which automates data integration steps by associating putative SVs with histone marks, TF binding sites, and genomic features, while accounting for both SV size distributions and co-occurring SNVs. As illustrated in Figure S1.B, CellVar provides a streamlined way to filter SVs within the scope of direct gene hits or to examine only enhancer candidates defined by published ChIP peak data (e.g., Encyclopedia of DNA Elements (ENCODE) neural crest stem cell H3K27ac peaks).

#### Creating OFV-DB

Lastly, all SV calls that passed the filtering process are formatted (normalized) into relational tables and saved into the MySQL database, called OFC DB along with the associated genomic features, genotype data and clinical variables. OFC DB can be easily queried to group SVs into distinct classes, such as those overlapping with genes, poised enhancer, active enhancer, or promoters as well as variants considered non-regulatory (Figure S1.C).

#### Statistical analysis

Our pipeline utilizes an ensemble approach to structural variant (SV) calling to maximize statistical power and optimize *in vivo* validation accuracy in human genomes. For Illumina paired-end data, individual reads retain a mapping quality (MAPQ) score, representing the alignment system’s probability estimation of a true genomic match. The ensemble framework evaluates these MAPQ metrics alongside insert-size distribution statistics to identify and validate true variant calls. Read depth callers like CNVnator, ERDS and CNVpytor use the read-depth statistics to make their calls. The details are available in the respective publications of the individual SV callers, and the experimental validation of the ensemble calling by FusorSV has been given in Becker et al. (2018).

### C. Iterative Feature Engineering with HITL for Pathway Extension

The next step is to predict which SVs may impact the dysregulation of craniogenesis, leading to nsOFC etiology. This pipeline is outlined in Figure S2, which highlights a two-tiered architectural framework for large-scale scientific data mining: (i) constructing a 3D OLAP-style data cube by integrating available SV data, and (ii) building a machine learning-based prediction system

#### 3D data cube generation incorporating multi-omics data and pathway data

The OFC-DB generated from our structural variant (SV) pipeline is augmented with two distinct public repositories relevant to craniogenesis: multi-omics datasets and functional pathway databases, including Reactome, KEGG and others (Figure S2.A). For instance, dataset GSE197513 (Yankee et al. (2023)) comprises bulk RNA-seq, single-cell RNA-seq (scRNA-seq), single-nuclei RNA-seq, and ChIP-seq modalities derived from human embryonic craniofacial tissues (collected at 4–8 weeks post-conception and Carnegie Stage 20) as well as cranial neural crest cells (CNCCs). These multi-omics data are preprocessed and dynamically mapped to the genomic features (GFs) overlapping our candidate SVs. This integration allows the system to score whether specific associated GFs are functionally influential during human craniogenesis.

**Figure S2.**
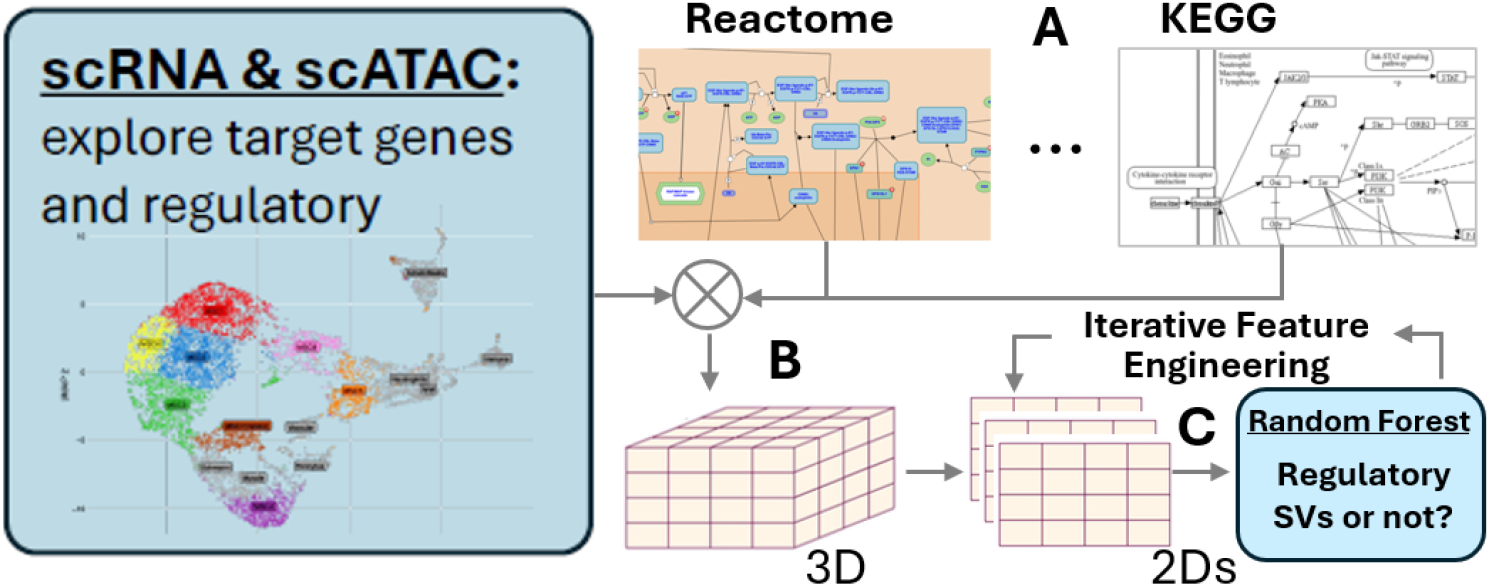
The overview of the workflow. A. integrating gene expression data with pertinent pathway data, B. Create 3D data structure for flexible data mining. C. Enabling prediction of novel SV genes through an iterative Human-in-the-Loop process.

Our scoring scheme also maps SV-associated genomic features (GFs) to specific developmental pathways. This mapping is critical to assessing whether the candidate features identified in a proband sample can explain disease etiology by disrupting key pathways curated for craniogenesis. While conventional methods rely on gene set enrichment analysis (GSEA) via Fisher’s exact test to identify involved pathways, we employ a deeper pathway topology analysis. Specifically, we map the sample-specific SV-GF gene lists onto distinct pathway “branches” or “routes” to gain a finer resolution of pathway disturbance. This route-based approach leverages rPAC (Joshi et al. (2022)), a locally developed method capable of revealing how multiple functional pathways are interconnected in a concerted manner.

The analytical outcomes from these multi-omics and developmental pathway data are modeled into 3D data cubes (Figure S2.B). These cubes are subsequently flattened into a series of 2D structures readily utilized for building machine learning models.

#### Human-in-the-Loop Refinement for Pathway Extension

The final stage of the workflow, outlined in Figure S2.C, implements an iterative feature engineering framework utilizing a Human-in-the-Loop (HITL) paradigm. Rather than relying on a static feature set or a fully automated wrapper method, this framework enables a human domain expert to dynamically manipulate and refine the feature vectors across successive training iterations. In each cycle, the algorithm evaluates individual information gain metrics for the underlying decision trees. The scientist reviews these outputs to manually add, remove, or modify features, leveraging contextual biological domain knowledge to test specific hypotheses. For example, the expert can evaluate the model’s response if ENCODE enhancer target prediction data from mesenchymal and neural crest cells are excluded; if SVs overlapping with exons are weighted more heavily; or if the specific weights of the cilia pathway are factored into the scoring system. The objective of this process is to determine which curated feature combination produces the highest overall classification accuracy, which is automatically computed through the established Random Forest model. Through this HITL framework, existing pathways are iteratively modified, ultimately producing an extended version of a developmental pathway crucial to craniogenesis—specifically, the epithelial-mesenchymal transition (EMT) pathway, which is well-documented as essential to palatogenesis. Our case studies in Supplemental Data Section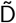 further illustrate the outcomes of this HITL approach applied to OFC-DB.

### D. Various Statistics

It is essential to provide a comprehensive description of the SV landscape in the WGS cohort study as part of quality control, cohort characterization, comparison with related studies and interpretation of downstream findings. We provide various summary statistics in this part of supplementary data.

**Figure S3.**
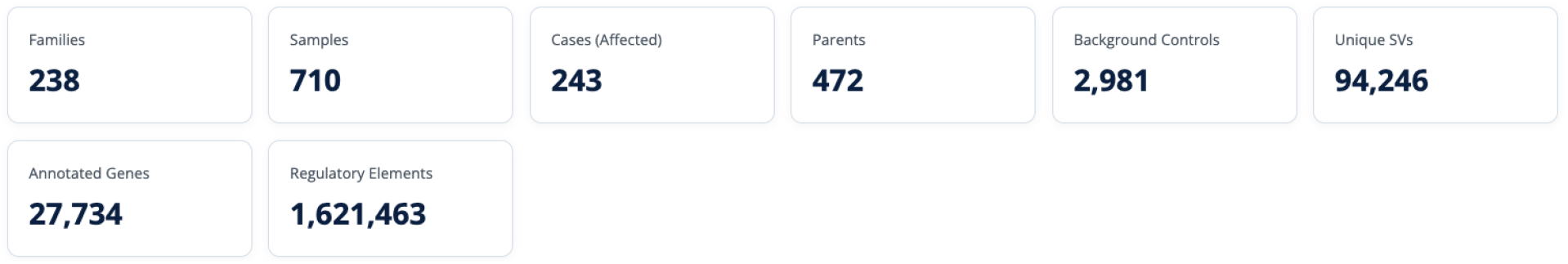
The African and Asian cohorts include total 241 families but due to inconsistencies found in sample annotations (father or mother missing, same gender parents), the total number of families analyzed reduced to 238. Regulatory elements here refer to exons, promoters, poised enhancers, active enhancers, and inhibitors.

**Figure S4.**
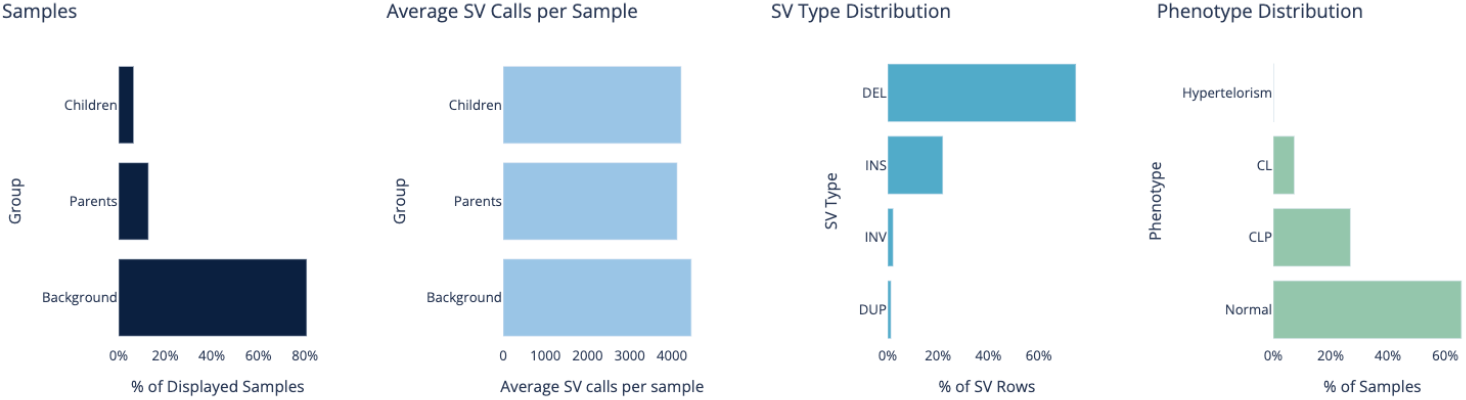
The sample group distribution clearly shows the dominant sample sizes of Parents and Background from the 1000 Genomes Phase 4 project. Noticeable is that the average SV calls per sample is uniform across the three different groups suggesting that the SV callling algorithms are not biased to any group, an important QC measure for the WGS data quality and the SV calling algorithms.

**Figure S5.**
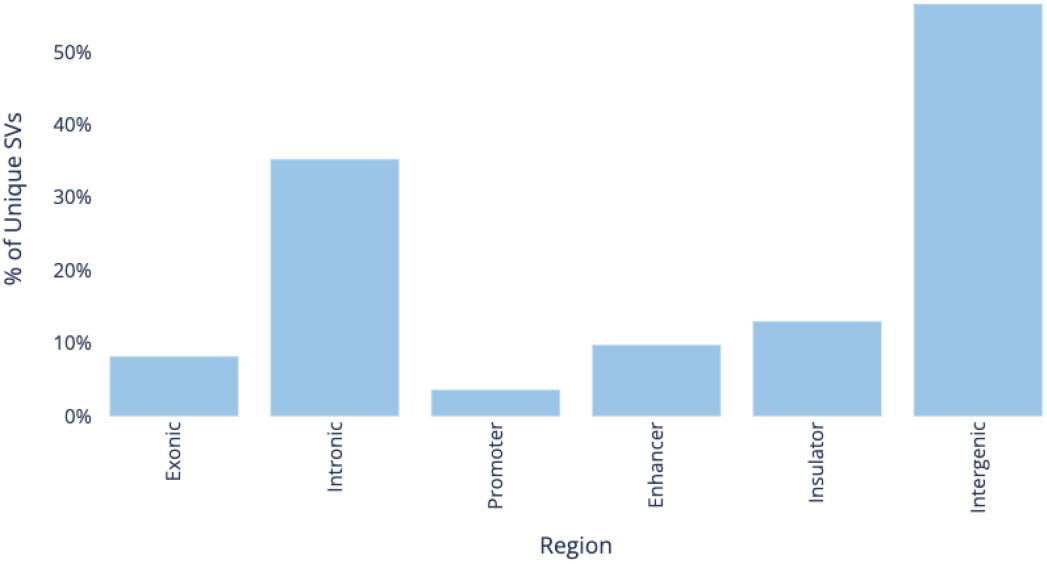
The regional distribution of SVs is consistent with patterns reported elsewhere, mapping predominantly to intergenic regions at 56.5% (53,290/94,246) followed by intronic regions at 35.3% (33,226/94,246). Notably, 3.6% (3,126/94,246) of SVs map to promoter regions.

**Figure S6.**
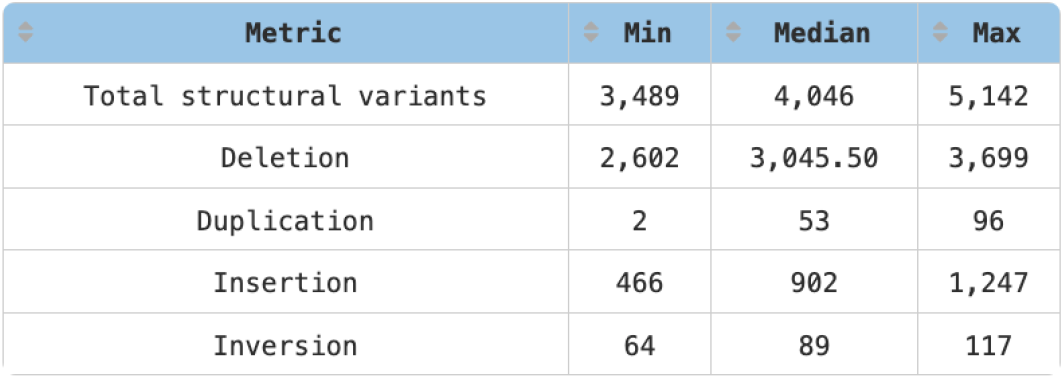
Consistent with published literature, the SV type distribution reveals that deletions occur most frequently and duplications least frequently. Furthermore, the minimal variance between minimum and maximum counts across all four categories reinforces the stringency of the SV calling algorithm’s quality matter.

**Figure S7.**
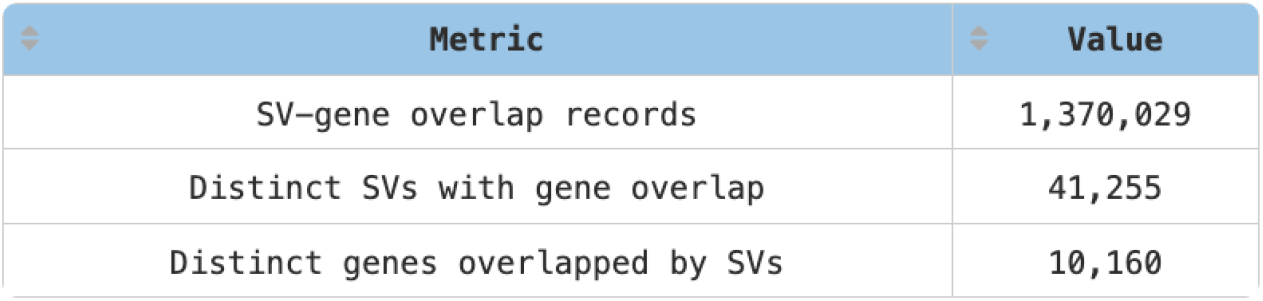
SV-gene overlap frequencies provide further characterization of the structural variant landscape. This analysis evaluates the data from two distinct angles: SV-centric (the number of unique SVs intersecting genes) and gene-centric (the number of unique genes intersecting SVs). From the gene-centric viewpoint, 36.6% (10,160/27,734) of all genes are overlapped by SVs. Combining both dimensions indicates an average density of approximately four SVs mapped per overlapped gene.

**Figure S8.**
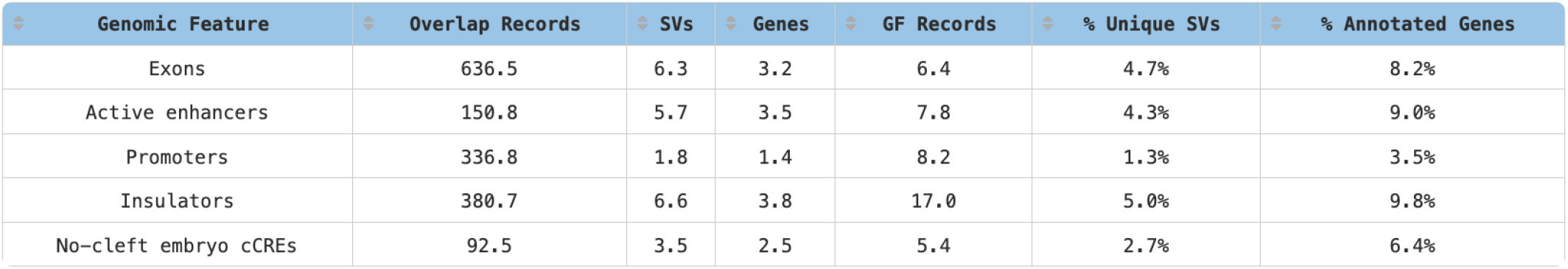
Per-sample analysis of SV overlaps across five distinct genomic features (GFs): exons, active enhancers, promoters, insulators, and no-cleft embryo cCREs. All listed counts represent cohort averages calculated by dividing total observations by the 710 samples in the dataset. GF annotations were derived from ENCODE data corresponding to mesenchymal and neural crest cells.

**Figure S9.**
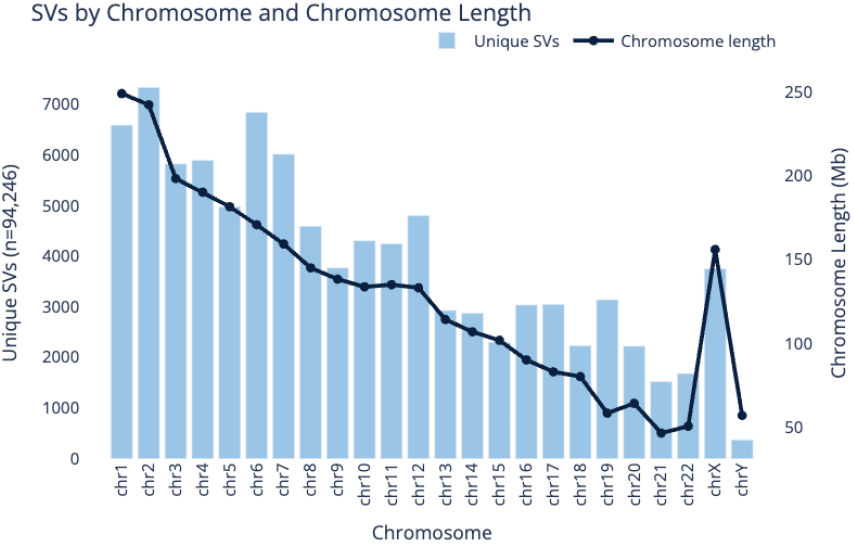
SV distribution across chromosomes follows a pattern similar to those in previous literature. The histogram is overlaid with chromosome lengths to show how SV frequency correlates with chromosome size. For instance, even though chr2 has a higher bar height than chr6, chr2’s larger size actually indicates a much higher frequency of SV events per unit length on chr6 relative to chr2. Similarly, the occurrence frequencies between chrX and chrY are roughly equivalent when adjusting for their physical sizes.

**Figure S10.**
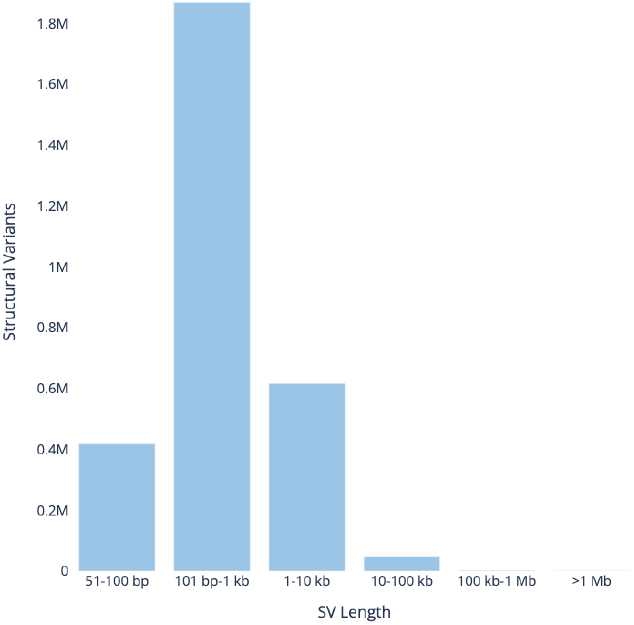
Size distribution of detected structural variants. The histogram that the majority of SVs fall within the 101 bp–1 kb range, whereas variants exceeding 1 Mb are rarely observed. Because SV calls were generated using Illumina short-read sequencing data, this length distribution should be interpreted within the context of short-read detection sensitivities and limitations.

### E. Pathway Extension Case Studies

#### Extending nsOFC Pathway Modeling

Our structural variant (SV) analysis identified 41 novel candidate genes, 13 of which share regulatory perturbations previously reported in the literature as SNV-associated risk genes (Supplementary Table 1). The objective of this section is to demonstrate how our HITL framework (discussed in Section C) extends the baseline nsOFC pathways documented in the literature, focusing specifically on six prioritized genes from these 41 candidates. Our long-term goal is to engage the craniofacial research community to collectively characterize the remaining 35 genes. We first present a summary matrix in Figure S13; the first three columns provide general gene annotations, while the remaining three columns summarize the outcomes of our in-depth functional analysis.

Next, to illustrate the impact of our HITL framework, we contrast the baseline developmental pathway (Figure S14) with the expanded network topology (Figure S15). The extended version incorporates additional gene nodes and regulatory edges identified during our analysis, mapping a more comprehensive regulatory landscape for craniogenesis.

**Figure S11.**
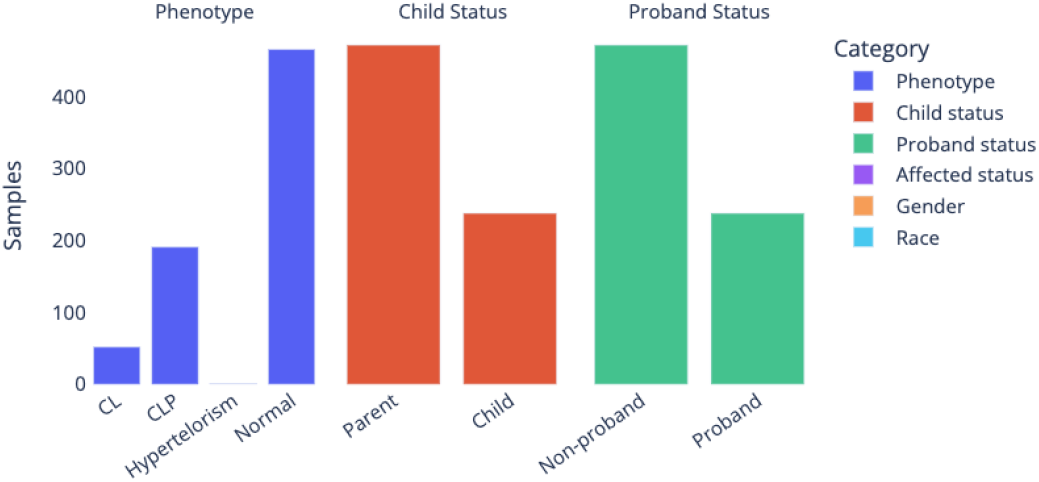
Cohort phenotypic and demographic breakdowns across six categories. The dataset includes closely balanced representation between African (n = 383) and Asian (n = 327) racial groups, enabling downstream analyses to detect whether specific SVs underlying nsOFC exhibit population-specific prevalence. Similarly, the balanced gender distribution provides sufficient statistical ground to investigate potential structural variants associated with sexual dimorphism.

**Figure S12.**
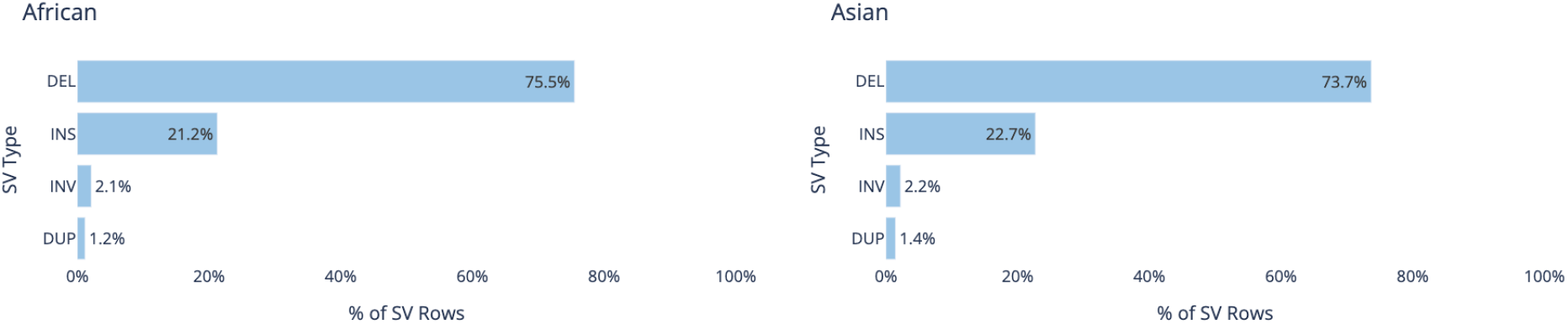
Stratified SV type composition for African and Asian study cohorts. Comparative analysis reveals a conserved distribution pattern across both groups. A marginal elevation in deletion events is noted within the African cohort, matching established literature benchmarks. Deletions (DEL) and insertions (INS) constitute the vast majority of events in both populations.

**Figure S13.**
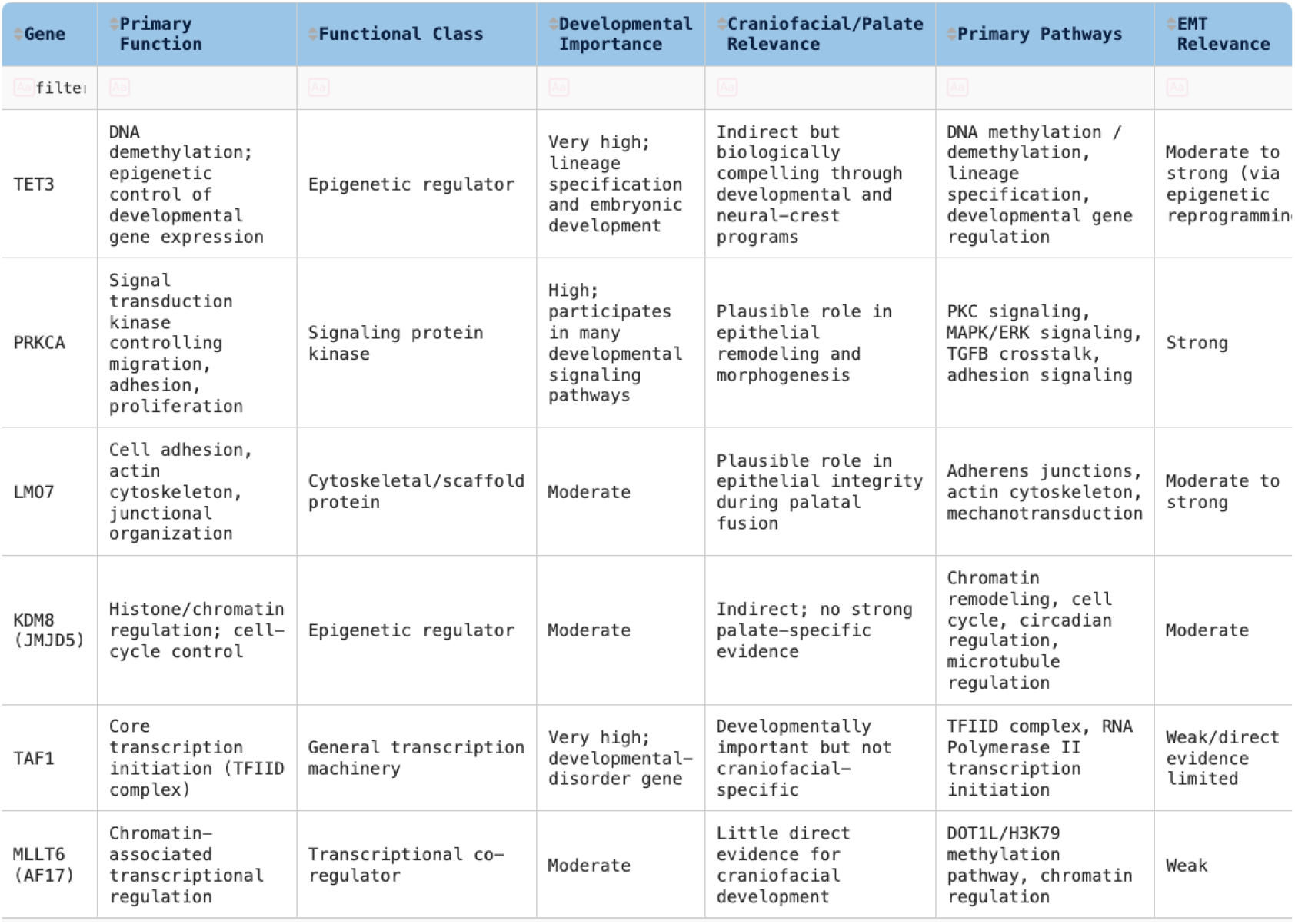
Functional annotations and pathway extension summary for six prioritized candidate genes. This table summarizes six genes harboring *de novo* or rare structural variants (SVs) identified in nsOFC probands. It highlights their regulatory mechanisms contributing to the nsOFC phenotype, evaluated across three primary dimensions: craniofacial/palatal relevance, primary developmental pathways, and epithelial-mesenchymal transition (EMT) relevance.

**Figure S14.**
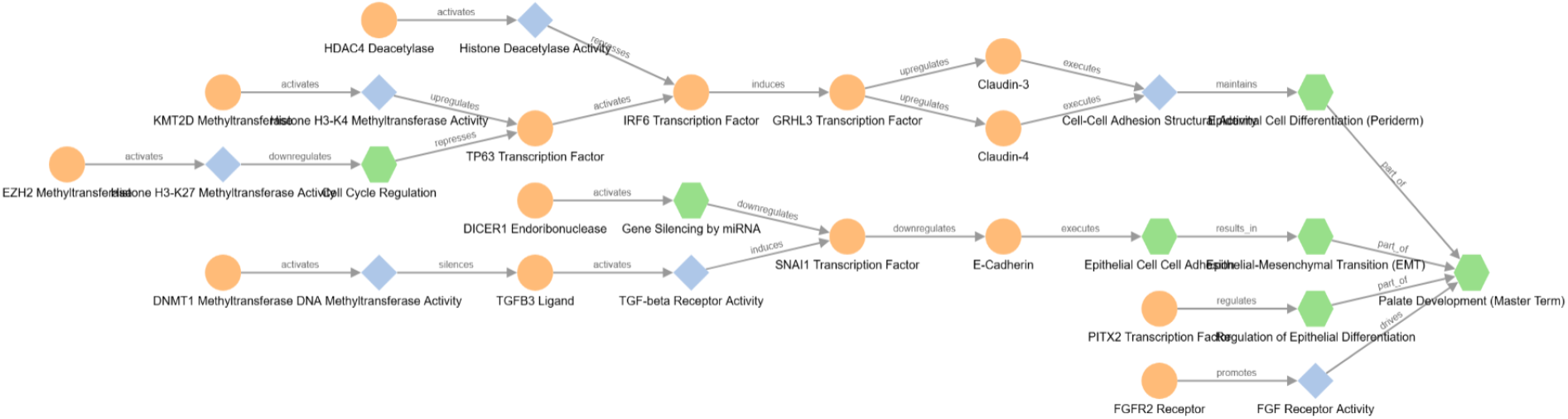
The baseline pathway diagram constructed by encoding regulatory perturbations previously reported in the literature. The gene and biological process nodes and relationship edges are mostly based on known SNV-associated risk genes for nsOFC.

**Figure S15.**
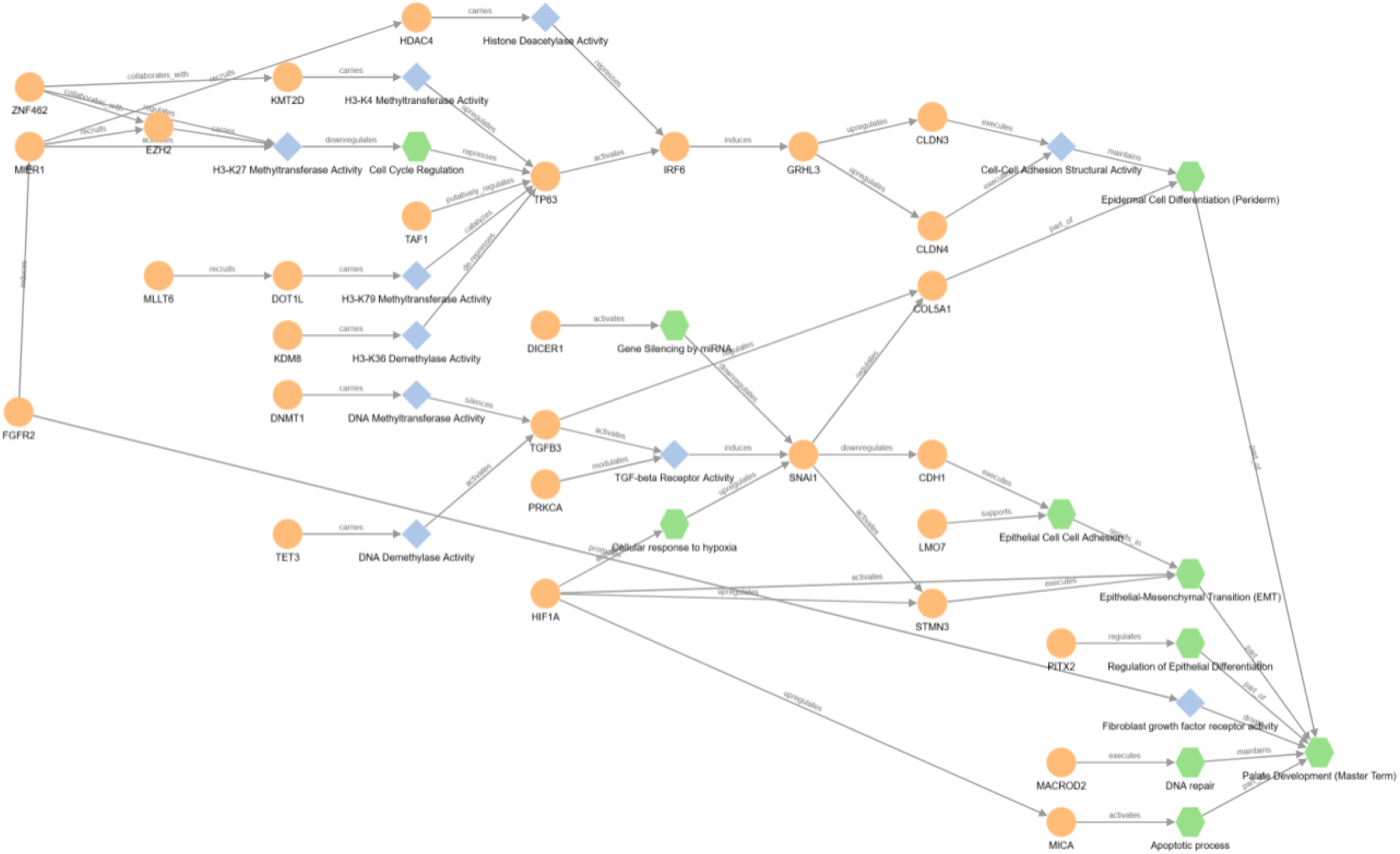
This extended pathway diagram provides a more comprehensive representation of genetic features and their relationships with epigenetic regulatory mechanisms associated with EMT. The newly curated regulatory mechanism identified through PRKCA processing is intentionally excluded to emphasize its distinct regulatory characteristics, which are illustrated separately in the stand-alone pathway diagram presented in Case Study 2.

#### Case study 1 - Coding exon disruption (TET3)

##### Literature

*TET3* is responsible for converting 5-methylcytosine (5mC) to 5-hydroxymethylcytosine (5hmC), which is responsible for initiating active DNA demethylation (Ketchum et al. (2024)). When *TET3* is compromised, downstream signaling required for neural crest cell differentiation and pharyngeal cartilage formation is impaired, which can lead to craniofacial malformations like cleft lip and palate. It has been reported that experimental loss of Tet2/3 disrupts *BMP*-dependent chondrocyte differentiation and pharyngeal cartilage formation (Wang et al. (2024)), suggesting that impaired *TET3* function could contribute to craniofacial malformations.

##### Pathway

Our analysis discovers the case(s) in rare frequency that demonstrates the oligogenic interaction involving *TET3*. The pathway diagram (Figure S16) shows which other well established genes (*SNAI1, CDH1, PITX2*) appear with *TET3* in our curated palatogenesis pathway diagram. The branches of the diagram interconnecting the selected genes show the involvement of developmental enhancers via the *TET-SNAIL4-BMP* axis.

**Figure S16.**
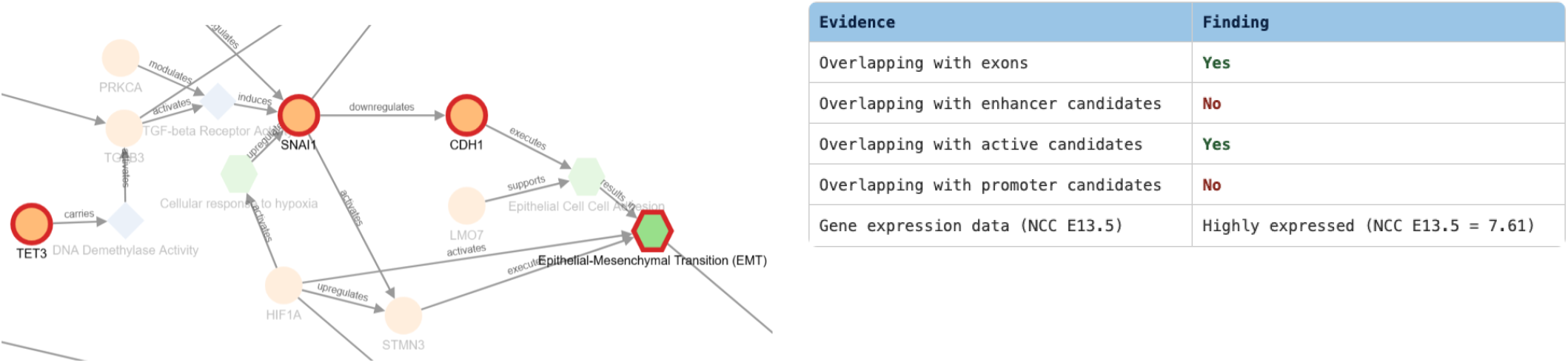
TET proteins are frequently found upstream of EMT programs because EMT requires large-scale epigenetic reprogramming. *TET* - mediated DNA demethylation can regulate expression of genes involved in: cell fate transitions, epithelial identity, migration, differentiation. It may not be an EMT transcription factor, but it can be labelled as EMT-enabling epigenetic regulator. Noticeable in the figure is that this gene is highly expressed in NCC E13.5.

#### Case study 2 - Promoter/TSS disruption (KDM8)

##### Literature

*KDM8* (*JMJD5*) is a chromatin-associated regulator implicated in control of chromatin state, cell proliferation, DNA damage responses, and cytoskeletal organization. Although it is not currently recognized as an established craniofacial developmental gene, its molecular functions could influence pathways relevant to development. *KDM8* contains a JmjC-domain oxygenase and was originally reported to demethylate H3K36me2, a histone modification associated with transcriptional regulation. Through modulation of H3K36 methylation and chromatin-associated protein recruitment, *KDM8* may influence the accessibility and expression of specific gene loci (Hsia et al. (2010), Ahmed et al. (2023)).

##### Pathway

Our analysis found KDM8’s promoter/TSS region overlapping with a deletion SV. Figure S17 depicts that KDM8 and DOT1L function as opposing biochemical forces on the epigenome, serving as an epigenetic eraser (demethylase) and an epigenetic writer (methyltransferase), respectively. Beyond modifying chromatin, KDM8 actively enters the cytoplasm or spindle apparatus to alter non-histone structures. It directly controls -tubulin acetylation, transforming physical microtubule architecture to assist the morphological softening and migration necessary for EMT. In contrast, DOT1L recognizes H3K79 within assembled nucleosomes and relies on extensive nucleosome interactions to position and methylate its target residue (zhang2019kdm8). While both drive the cellular plastic changes of the epithelial-to-mesenchymal transition (EMT), they differ entirely in their catalytic mechanisms, targeted histone marks, and non-histone structural actions.

**Figure S17.**
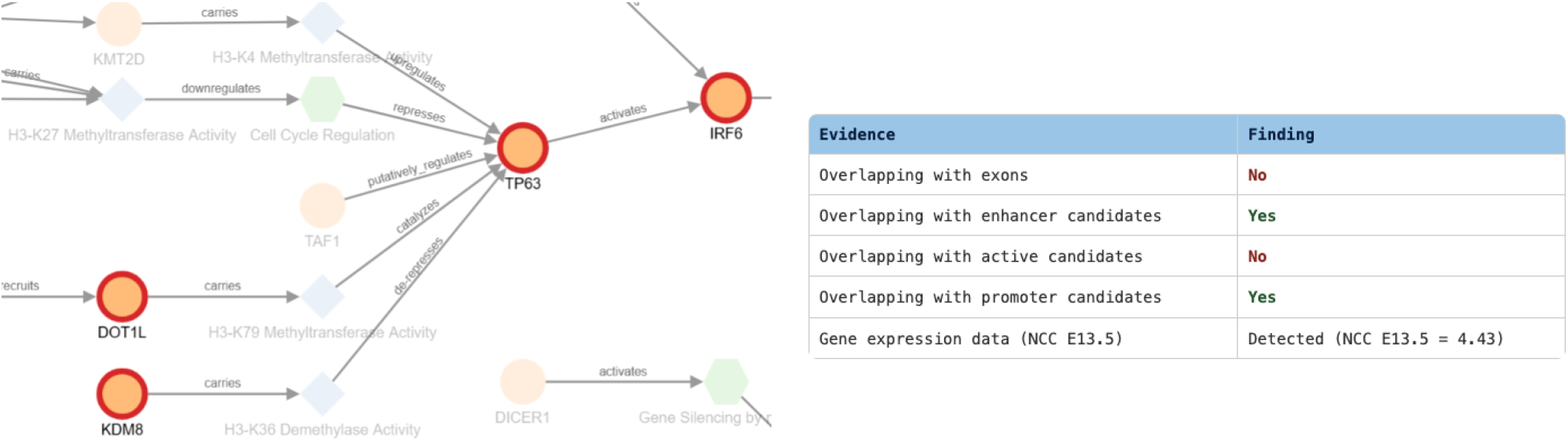
The pathway diagram highlights the complementary epigenetic regulatory functions of *DOT1L* and *KDM8*, with *DOT1L* mediating H3K79 methylation and *KDM8* contributing to regulation of H3K36 methylation. Disruption of the *KDM8* promoter/TSS region is hypothesized to alter chromatin regulation and downstream transcriptional programs, potentially affecting palatogenesis through pathways that converge on the established *TP63-IRF6-GRHL3* regulatory axis. This figure shows that KDM8 is detected in NCC E13.5.

#### Case study 3 - Intergenic distal enhancer disruption (MLLT6)

##### Literature

*MLLT6* is a chromatin-associated transcriptional regulator that interacts with histone-modifying machinery. One of its best-characterized functions is interaction with the *DOT1L/DOT1A* H3K79 methyltransferase complex, where *MLLT6* competes with *AF9* and can alter *DOT1L* localization and H3K79 methylation levels (Reisenauer et al. (2009)).

##### Pathway

In Figure S18, *MLLT6* is positioned upstream of *DOT1L* as a chromatin-associated regulator that may facilitate *DOT1L*-dependent H3K79 methylation. This epigenetic module is hypothesized to support transcriptional programs required for epithelial differentiation and integrity, thereby influencing developmental pathways that converge on the *TP63-IRF6-GRHL3* network during palatogenesis.

**Figure S18.**
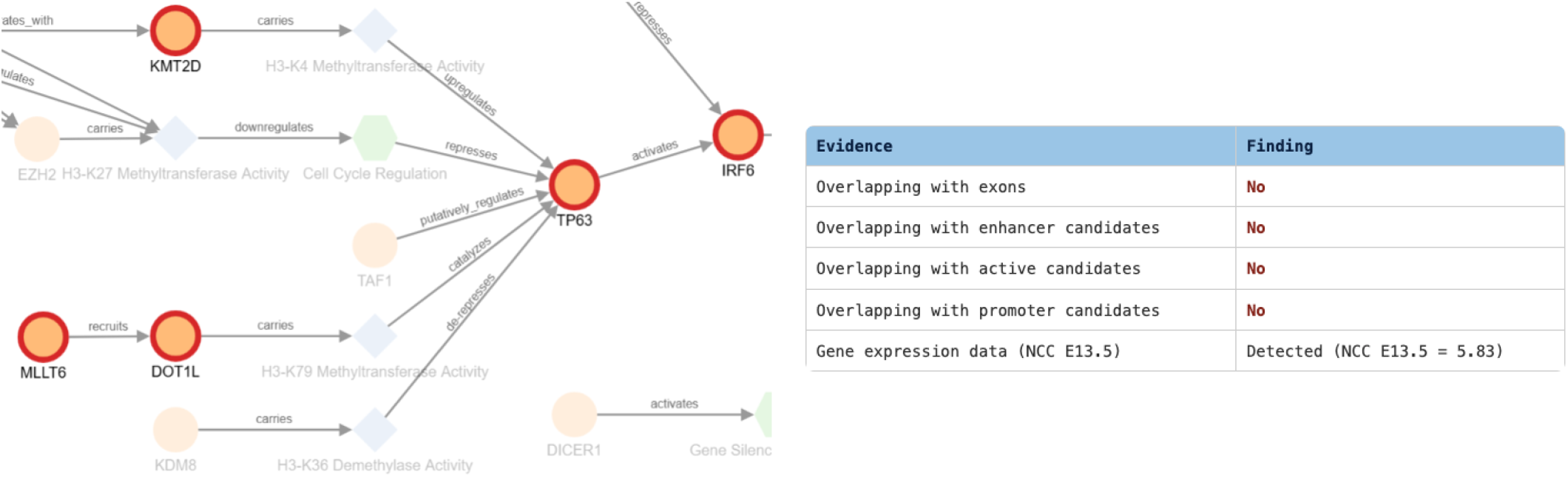
The pathway proposes that both the *KMT2D* and *MLLT6-DOT1L* modules act as positive regulators of the *TP63* developmental program. While *KMT2D* contributes enhancer activation and *DOT1L* contributes H3K79 methylation-associated transcriptional activity, both pathways are predicted to converge on maintenance of *TP63-*dependent epithelial differentiation, including the downstream *IRF6-GRHL3* axis.

#### Case study 4 - Intron enhancer disruption (LMO7)

##### Literature

In several cancer models, loss or dysregulation of LMO7 can alter cell morphology, migration, invasion, and EMT-associated programs Zhen et al. (2022); Zeng et al. (2024); Liu et al. (2021). In addition, *LMO7* has been implicated in the organization and remodeling of embryonic epithelial tissues through its role in apical junctional complexes and actomyosin-dependent epithelial morphogenesis (Matsuda et al. (2022)).

##### Pathway

In Figure S19, *LMO7* is positioned as a contributor to epithelial cell-cell adhesion. Adherens junctions depend on both cell-cell recognition/adhesion and mechanical coupling to the actin cytoskeleton, with *LMO7* contributing primarily to the latter. Through interactions with afadin and -actinin, *LMO7* helps organize the actin network associated with adherens junctions, thereby supporting the stabilization and maintenance of epithelial cell-cell contacts.

**Figure S19.**
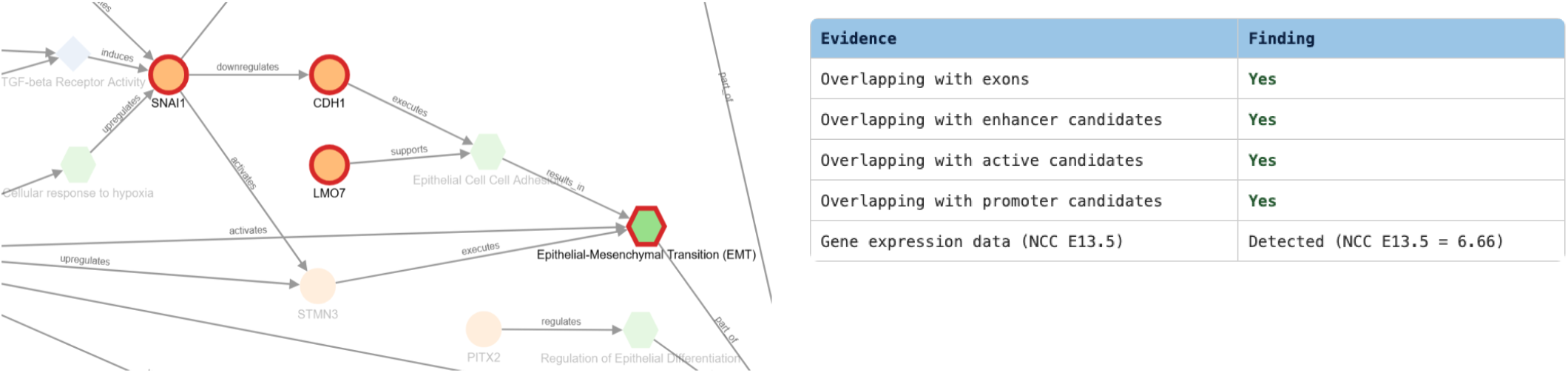
The extended pathway diagram shows how *LMO7* is placed. It supports Epithelial Cell-Cell Adhesion leading to EMT but in parallel to the known regulatory path *SNAI1* - *CDH1* - Epithelial Cell-Cell Adhesion - Epithelial-Mesenchymal Transition (EMT).

#### Cast study 5 - Intergenic insulator/CTCF disruption (TAF1)

##### Literature

Animal models show that disrupting the TAF complex leads directly to craniofacial hypoplasia. A shift in *TAF1* expression in cranial neural crest cells would alter downstream skeletal patterning, mimicking phenotypes seen in *TAF1* intellectual disability syndromes (such as distinct facial dysmorphism). Migrating neural crest cells require precision control of their transcriptomes to successfully navigate the embryo. Altered *TAF1* dosage shifts the expression profiles of core genes, potentially stalling cell migration (Leid et al. (2023), Hurst et al. (2018), Gudmundsson et al. (2019))

##### Pathway

*TAF1* is the largest subunit of the TFIID complex, serving as a major scaffold and regulatory component within *TFIID*. Our SV analysis discovered a 100-bp deletion structural variant disrupting an insulator 13.8 kb upstream of *TAF1* as potentially breaking a non-coding structural component like an insulator changes the physical 3D architecture of the DNA. In Figure S20, *TAF1* is linked to *TP63* with the relationship “putatively regulate” based on these known facts: *TP63* is a transcription factor controlling epithelial and periderm gene programs; *TP63* target genes must be transcribed through the general RNA polymerase II machinery; *TFIID* (containing *TAF1*) is a key component of that machinery; Thus, altered *TAF1* function could impair expression of *TP63* -regulated genes.

**Figure S20.**
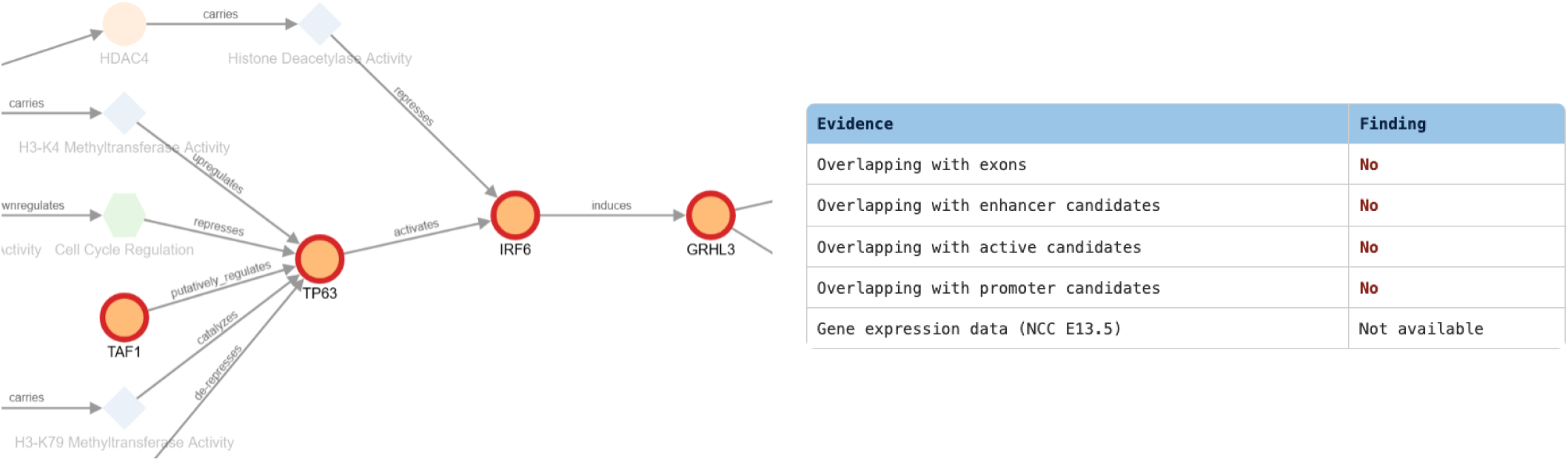
Our pathway extension introduces the relationship “putatively regulates” to distinguish predicted interactions from established ones. In this example, *TAF1* is predicted to regulate *TP63*, whereas the downstream *TP63* -*IRF6* -*GRHL3* regulatory axis is already established in the baseline pathway diagram.

#### Case study 6 - Intronic insulator/CTCF disruption (PRKCA)

##### Literature

*PRKCA* is essential for Neural Crest Migration. Unlike structural cilia genes, *PRKCA* is highly active and structurally vital in migrating neural crest cells. Epithelial-to-mesenchymal transition (EMT) and directed cell migration require constant changes to the cell skeleton. *PRKCA* signaling regulates the focal adhesions and actin dynamics that allow cranial and trunk neural crest cells to break away and travel through the embryo (Patel et al. (2020))

##### Pathway

Our SV analysis revealed a 100-bp deletion inside an intron of *PRKCA* that overlaps a known 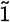-kb mesenchymal CTCF/insulator. It can be thought of as a small regulatory/chromatin-architecture SV affecting the *PRKCA* locus, rather than an intronic loss-of-function variant. *PRKCA* encodes protein kinase C, a signaling kinase involved in proliferation, differentiation, adhesion, and cytoskeletal regulation, processes that are central to craniofacial morphogenesis and orofacial cleft development (Ji et al. (2020)).

In Figure S21, *PRKCA* is depicted as a regulatory brake that modulates cellular responses to incoming signals, whereas *MKKS* and *DNAJC5* act as structural components required for the assembly and function of the primary cilium, where *PRKCA* operates as a localized signaling kinase (Reddy (2005)). In addition, *PRKCA* is placed as a direct modulator of the SHH Signaling node (GO:0007224) (Reddy (2005), Mukhopadhyay and Rohatgi (2014)).

**Figure S21.**
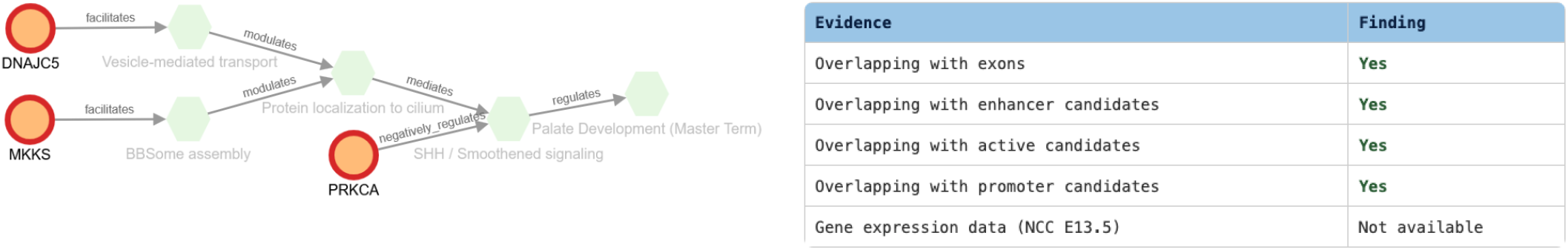
It shows how Intronic insulator/CTCF disruption of PRKCA could tip the balance between downstream regulatory activities of DNAJCS and MKKS implicated in Protein location to cilium and eventually lead to dysregulation of Palate Development.

